# DRAMS: A Tool to Detect and Re-Align Mixed-up Samples for Integrative Studies of Multi-omics Data

**DOI:** 10.1101/831537

**Authors:** Yi Jiang, Gina Giase, Kay Grennan, Annie W. Shieh, Yan Xia, Lide Han, Quan Wang, Qiang Wei, Rui Chen, Sihan Liu, Kevin P. White, Chao Chen, Bingshan Li, Chunyu Liu

**Affiliations:** Center for Medical Genetics & Hunan Key Laboratory of Medical Genetics, School of Life Sciences, Central South University, Changsha, Hunan, China; Department of Molecular Physiology & Biophysics, Vanderbilt University, Nashville, TN 37212, USA; Vanderbilt Genetics Institute, Vanderbilt University Medical Center, Nashville, TN, USA; School of Public Health, University of Illinois at Chicago, Chicago, IL, USA; Department of Psychiatry, SUNY Upstate Medical University, Syracuse, NY, USA; Institute for Genomics and Systems Biology, Department of Human Genetics, University of Chicago, Chicago, IL, USA; Tempus Labs Inc, Chicago, IL, USA; National Clinical Research Center for Geriatric Disorders, Xiangya Hospital, Central South University, Changsha, Hunan, China; School of Psychology, Shaanxi Normal University, Xi’an, Shaanxi, China

## Abstract

Studies of complex disorders benefit from integrative analyses of multiple omics data. Yet, sample mix-ups frequently occur in multi-omics studies, weakening statistical power and risking false findings. Accurately aligning sample information, genotype, and corresponding omics data is critical for integrative analyses. We developed DRAMS (https://github.com/Yi-Jiang/DRAMS) to Detect and Re-Align Mixed-up Samples to address the sample mix-up problem. It uses a logistic regression model followed by a modified topological sorting algorithm to identify the potential true IDs based on data relationships of multi-omics. According to tests using simulated data, the more types of omics data used or the smaller the proportion of mix-ups, the better that DRAMS performs. Applying DRAMS to real data from the PsychENCODE BrainGVEX project, we detected and corrected 201 (12.5% of total data generated) mix-ups. Of the 21 mix-ups involving errors of racial identity, DRAMS re-assigned all samples to the correct racial cluster in the 1000 Genomes project. In doing so, quantitative trait loci (QTL) (FDR<0.01) increased by an average of 1.62-fold. The use of DRAMS in multi-omics studies will strengthen statistical power of the study and improve quality of the results. Even though very limited studies have multi-omics data in place, we expect such data will increase quickly with the needs of DRAMS.

**Author summary:** Sample mix-up happens inevitably during sample collection, processing, and data management. It leads to reduced statistical power and sometimes false findings. It is of great importance to correct mixed-up samples before conducting any downstream analyses. We developed DRAMS to detect and re-align mixed-up samples in multi-omics studies. The basic idea of DRAMS is to align the data and labels for each sample leveraging the genetic information of multi-omics data. DRAMS corrects sample IDs following a two-step strategy. At first, it estimates pairwise genetic relatedness among all the data generated from all the individuals. Because the different data generated from the same individual should share the same genetics, we can cluster all the highly related data and consider that the data from one cluster have only one potential ID. Then, we used a “majority vote” strategy to infer the potential ID for individuals in each cluster. Other information, such as match of genetics-based and reported sexes, omics priorority, etc., were also used to direct identifying the potential IDs. It has been proved that DRAMS performs very well in both simulation and PsychENCODE BrainGVEX multi-omics data.

## Introduction

Investigation of complex traits and disorders can use multiple omics data to systematically explore regulatory networks and causal relationships. Sample mix-ups can occur in omics experiments during sample collection, handling, genotyping, and data management. As the number of datasets to be integrated increase, the likelihood of error also multiplies. Sample mix-ups reduce statistical power and generate false findings. Not only is the detection and re-alignment of errors in data identifications (IDs) critical to ensuring accurate findings in integrative studies, such corrections can increase statistical power thus the number of positive findings [1].

For multi-omics data, the sample re-alignment procedure can be generally divided into two steps: first, to estimate genetic relatedness among the data of different omics and cluster together all the data of the same individual; then, to assign potential IDs for each data cluster. It is well-known that genetic information from the same individual should be identical regardless of the omics from which it originated. Using genotype data as a mediator, data originated from the same individual can be clustered together.

Several tools have been developed to estimate genetic relatedness for multi-omics data in many different ways, such as genotype concordance [2, 3], correlation of different quantifications [1, 2], correlation of variant allele fractions [4], concordance of sequencing reads [4], *etc*. The various methods made it possible to compare different data types, such as DNA sequencing, RNA sequencing, SNP array, *etc*. However, these tools mainly focused on implementing the first step of sample re-alignment, that is, to estimate genetic relatedness among multi-omics data. None of the existing tools has a systematic solution on determining the potential IDs for each data.

It is certain that, after clustering the highly related data, some data ID can be empirically corrected based on the “majority vote” strategy [1, 2, 5]. However, without a systematic solution, it only works for small data of low-dimension, difficult to scale up, and the results lack statistics-based confidence. As multi-omics data of more data dimension are expected in the near future, it is much more challenging to identify the potential ID. Especially when more than one data are labeled by mistake for a single individual. Actually, taking full advantage of information from data of more omics types makes data ID correction more accurate, turning a challenge into an opportunity.

Here, we described DRAMS, a tool to <u>D</u>etect and <u>R</u>e-<u>A</u>lign <u>M</u>ixed-up <u>S</u>amples that leverages sample relationships in multi-omics data by directly comparing genotype data. DRAMS also uses a logistic regression model followed by a modified topological sorting algorithm to systematically re-align misclassifications. This tool integrates sample relationships among different omics types, concordance rates of genetics-based sexes and reported sexes, etc. With this design, DRAMS can be applied to as many omics types as possible. Using both simulated data and real data, we proved the accuracy and power of DRAMS in studies involving multi-omics data.

## Results

### Design of DRAMS

The goal of DRAMS is to detect and re-align mix-ups based on the grounds that all omics data originating from the same individual should match genotypes. DRAMS operates on a two-step process: first, we ensure that all omics data of the same samples were cluster together by their genotypic relatedness; after that, we find out the potential true ID of each data cluster (Fig 1).

**Fig 1.**
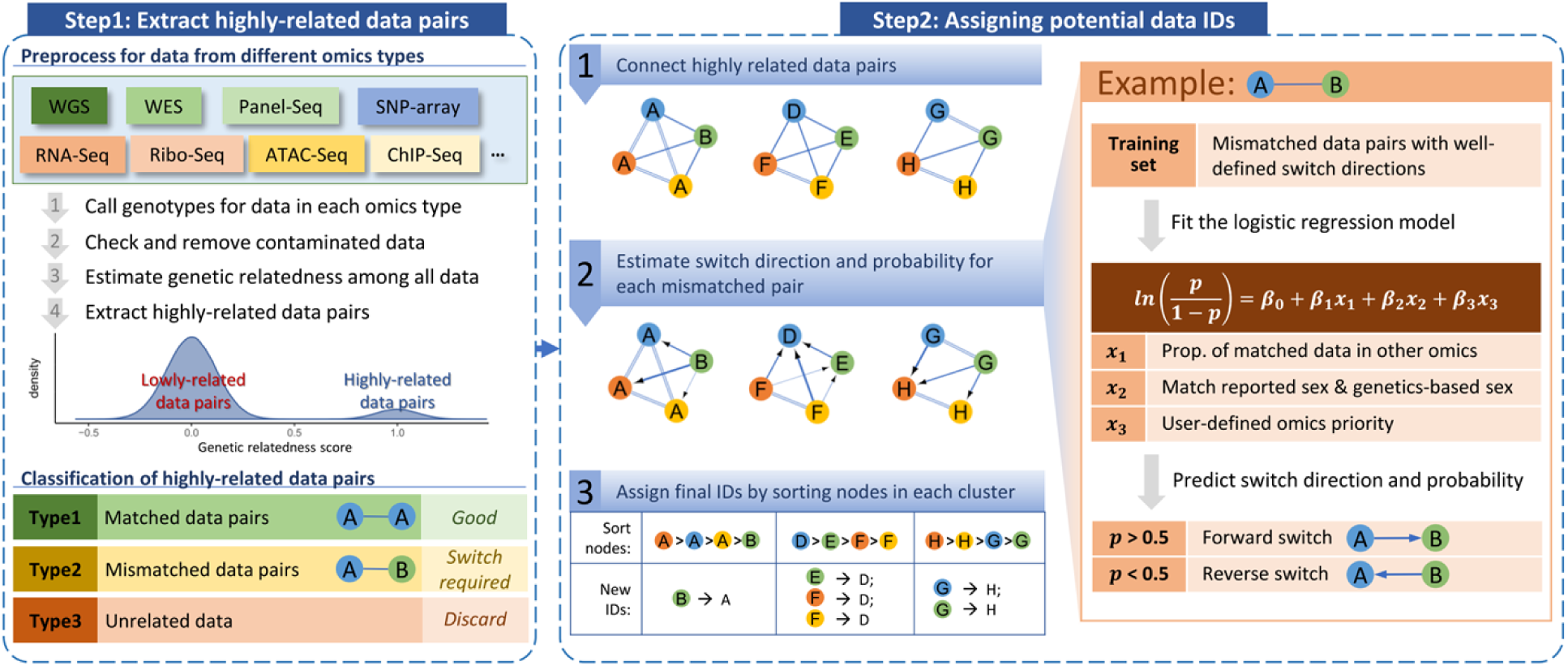
Illustration of key steps. The workflow has two basic steps: 1) extracting highly related data pairs; and 2) assigning potential data IDs. The first step calls genotypes for data from each omics types. DRAMS then estimates genetic relatedness by comparing genotypes among all available omics data. Any omics type that has the potential to call genotypes can be used as input. Highly related data pairs are extracted based on the distribution of relatedness scores and connected to create multiple clusters. In the second step, each node represents a data in the cluster. The text in each node represents the data ID. Different colors represent different omics types. The parallel line connects *matched* data; whereas, the singular line connects *mismatched* data. After applying a logistic regression model, DRAMS tool estimates switch directions and probabilities for each mismatched data pair. The arrow denotes possible switch direction. The thickness of the line weight correlates to the degree of switch probability. The final IDs for the data in each cluster are determined by sorting the nodes.

For the first step is to build highly related data pairs. To accomplish this, we call genotypes from each omic dataset and estimate genetic relatedness among these data. When clustering omics data according to the highly-related data pairs, we can classify relationships of data into three types. The first type contains data pairs that have the same individual IDs (matched pairs). These are the least likely to be mis-assigned. The second type contains data pairs with different individual IDs (mismatched pairs), in which some individual IDs may have been swapped. We also have some data that are unrelated to any other. We put them into the third type, which will be discarded since they are unassignable.

For the second step, we connect all the highly related data pairs and produce multiple independent clusters. Each node within a cluster represents one data point with each edge connecting a highly related data pair. Then, we use a multi-step, combined knowledge-based and statistical approach to search for the potential IDs in each of the node clusters, which contains both matched and mismatched data pairs. To estimate which data ID from the mismatched data pair was more likely a correct ID, we first use a logistic regression model. The model estimates the direction and probability based on three pieces of information: 1) data relationships among multiple omics types –data matching a greater number of omics sets are more likely to represent the true ID than those matching only a few; 2) when the reported sex of the data agrees with the genetics-based sex, it is more likely to be accurate than not; 3) user-defined priority of omics data: the user’s confidence in the correctness of each omics type is documented as ranks. We manually identified 44 high-confidence mismatched data pairs with well-defined switch directions from the PsychENCODE BrainGVEX project[6] and used them to train the logistic regression model and establish parameters. **(Methods)**. In the last step, we determine the final ID for all omics data points within each cluster. We sort all data points in each cluster using a modified topological sorting algorithm that is based on the switch probabilities obtained from the logistic regression above **(Methods)**. The data ID with the highest value in each cluster will be selected as the final ID. In this study, we used simulation data and the PsychENCODE BrainGVEX data to evaluate the performance of DRAMS.

### Performance of DRAMS using simulation data

To test the performance of DRAMS on correcting data IDs, we generated multiple highly related data pair datasets with a few samples (ranging from 5% to 30%) being randomly shuffled to simulate sample mix up. The variable parameters for the simulation data include a range of sample sizes, numbers of omics types, and percentages of mix-ups (**Methods**). We found that when a larger proportion of samples are mixed up, fewer samples can be successfully corrected (Fig 2, S3 Table). Taking datasets with five omics types and a sample size of 300 as examples, when 5% of the samples are mis-assigned, all samples can be successfully corrected; when 30% of the samples are mis-assigned we can successfully correct an average of 92.4% (SD: 2.15%) of errors (Fig 2c).

**Fig 2.**
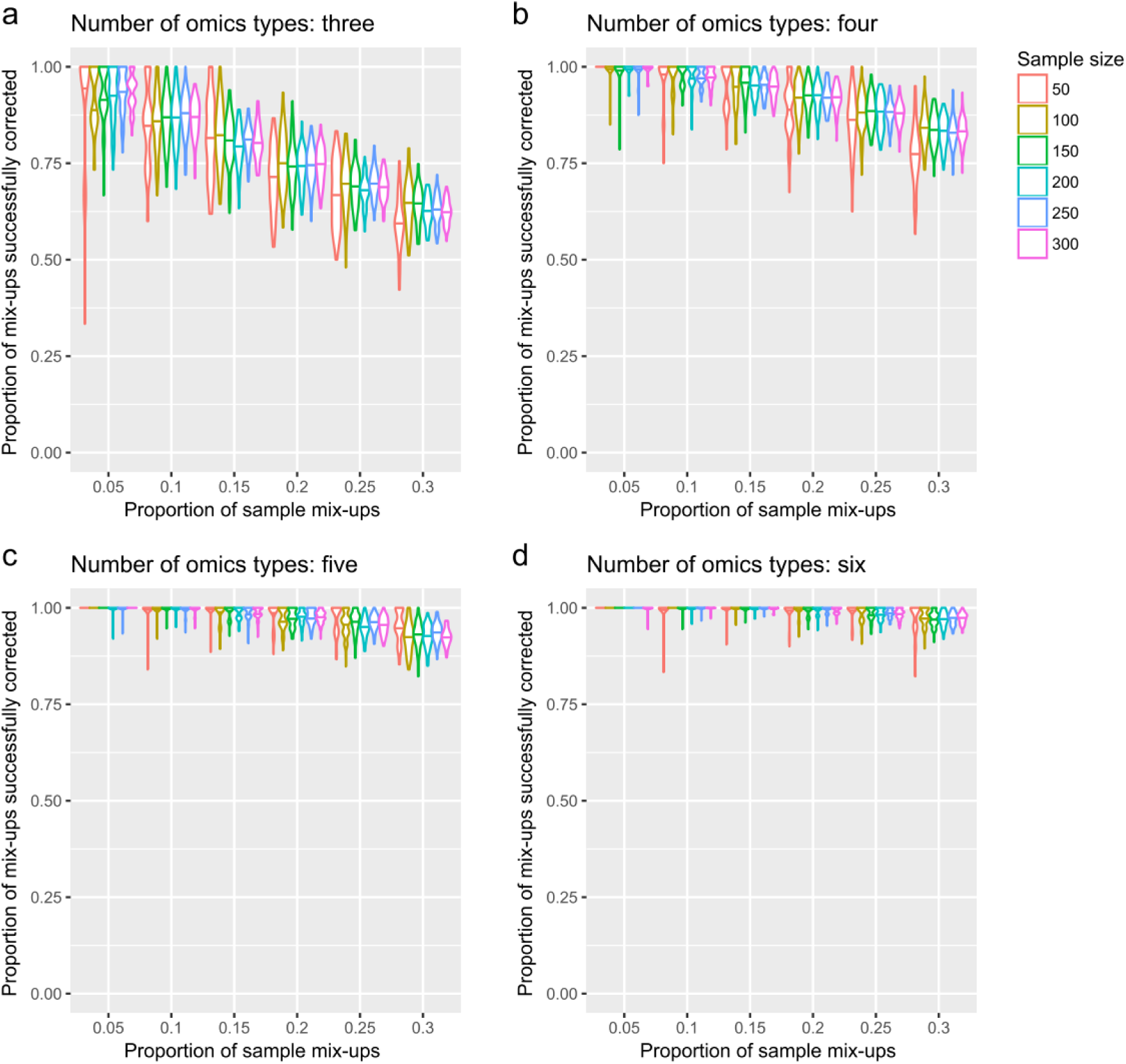
Performance of DRAMS in simulation data. We simulated sample mix-ups and used DRAMS to correct data IDs. The simulation data includes a range of sample sizes (50, 100, 150, 200, 250, and 300, each with 50% females and 50% males) and a range of omics types (3, 4, 5, and 6 for figure a, b, c, and d, respectively). To simulate sample mix-ups, we randomly shuffled a gradient proportion of data for each omics type: 5%, 10%, 15%, 20%, 25%, and 30%. The simulation process was repeated 100 times.

The number of omics types involved also greatly influences the performance of DRAMS. The more omics types we have, the better the chance that we can recover the true identity of each erroneous data. Taking datasets with a sample size of 300 and 15% of the samples mixed up as examples, if we have three omics types (the minimum number of omics types required for DRAMS input), an average of 80.5% (SD: 4.50%) of mix-ups can be successfully corrected (Fig 2a). If six omics types are used, an average of 99.7% (SD: 0.626%) mix-ups can be successfully corrected (Fig 2d).

Sample size is also important to the performance of DRAMS. Yet, sample size has no significant influence on DRAMS performance when proportions of sample mix-ups and number of omics used remain consistent (Pearson correlation between sample size and average proportions of successfully corrected mix-ups: 0.00901; P value: 0.915). However, larger sample sizes seem to stabilize correction results. As sample size increases, the standard deviation of the proportions of successfully corrected mix-ups decreased (Pearson correlation: −0.351; P value: 1.62 × 10-5).

### Performance of DRAMS using real data from the PsychENCODE BrainGVEX project

#### Data summary

The PsychENCODE BrainGVEX project[6] generated six types of omics data (S1 Table), including low-depth Whole Genome Sequencing (WGS), RNA sequencing (RNA-Seq), Assay for Transposase-Accessible Chromatin using Sequencing (ATAC-Seq), Ribosome Sequencing (Ribo-Seq), and SNP array data from two platforms, including Affymetrix 5.0 450K (Affymetrix) and Psych v1.1 beadchips (PsychChip). We called a total of 19,242,755 SNPs from WGS data on autosomes, 17,786,350 SNPs from RNA-Seq data, 10,571,742 SNPs from ATAC-Seq data, 156,354 SNPs from Ribo-Seq data, 10,891,109 SNPs from Affymetrix data, and 13,589,867 SNPs from PsychChip data. We used two methods to check sample contamination: using VerifyBamID[7], and calculating heterozygous rates (S1 Fig). We defined the samples with both FREEMIX >0.3 and heterozygous rates >0.3 as contaminated samples. We removed the sample “2015-916” in WGS from the subsequent analyses for being contaminated.

#### Check sample alignment based on genetics-based sex

We checked sample alignment by comparing genetics-based sexes with reported sexes for data of WGS, PsychChip, ATAC-Seq, and RNA-Seq. Based on X chromosome heterozygosity and Y chromosome call rate, we calculated an F-value (A Plink-derived metric to distinguish males and females. Details in Methods) for each data in each omics type. Then, the genetics-based sexes were inferred according to the distribution of F-values (called SNP-inferred sexes). By comparing reported sex with SNP-inferred sex, we identified a total of 74 data with mismatched sexes and 1174 data with matched sexes, indicating that some samples might have been mixed-up (Table 1). For the 426 samples in RNA-Seq, only three samples were identified as having mismatched sexes, indicating that this RNA-Seq data may have an overall good quality in terms of sample matching. For Ribo-Seq data, we did not estimate SNP-inferred sexes as the threshold to separate males and females, since Ribo-Seq data cannot be defined based on the distribution of F-values. Neither did we estimate SNP-inferred sexes for Affymetrix 5.0 SNP array data since no genotype on sex chromosomes were available. As complementary evidence, we also used *XIST* gene expression level (XIST-inferred sexes) to infer genetics-based sexes for RNA-Seq and Ribo-Seq data **(Methods)**.

**Table 1.**
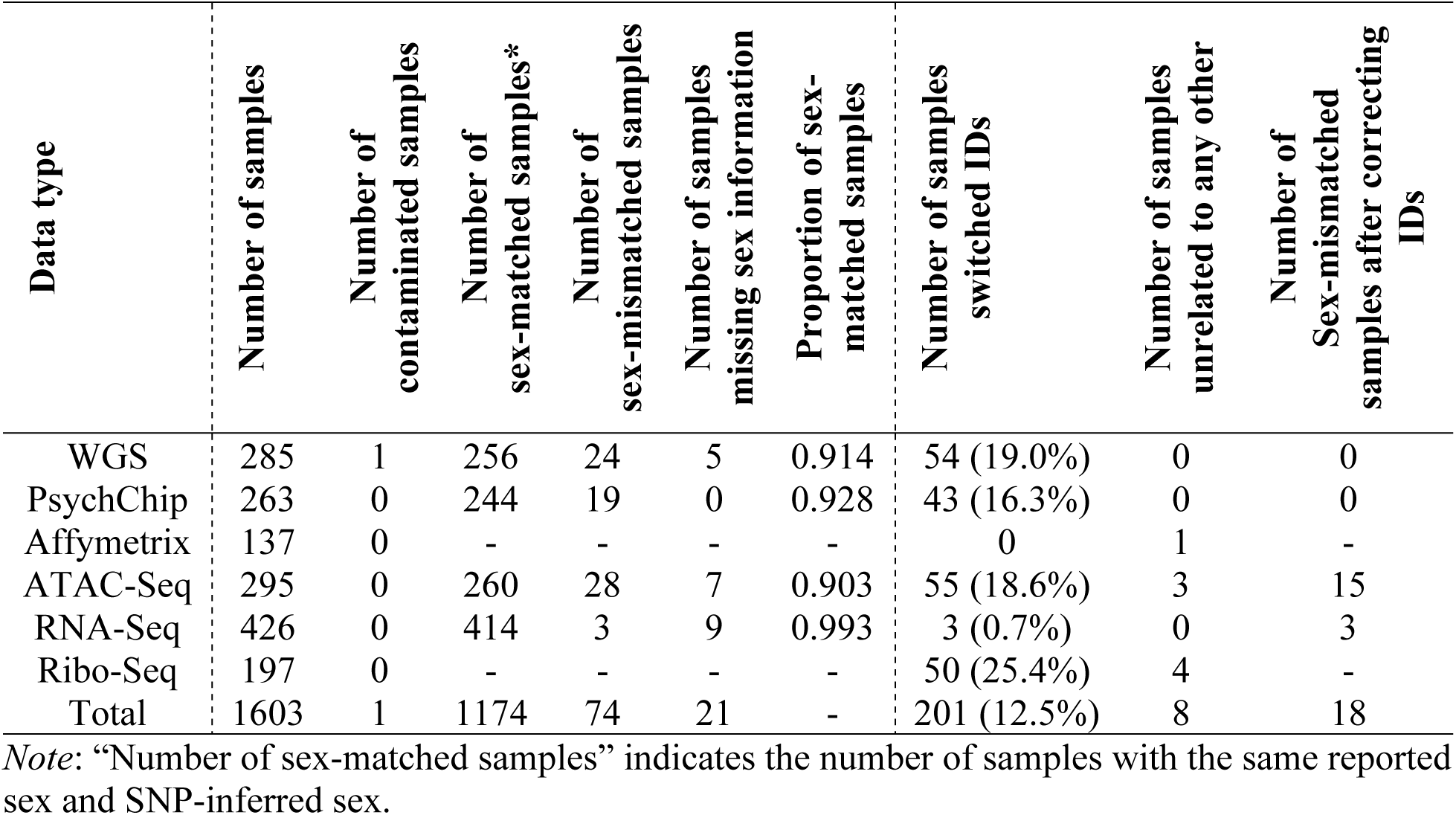
Summary of samples and sample corrections in BrainGVEX

| Data type | Number of samples | Number of contaminated samples | Number of sex-matched samples* | Number of sex-mismatched samples | Number of samples missing sex information | Proportion of sex-matched samples | Number of samples switched IDs | Number of samples unrelated to any other | Number of Sex-mismatched samples after correcting IDs |
| --- | --- | --- | --- | --- | --- | --- | --- | --- | --- |
| WGS | 285 | 1 | 256 | 24 | 5 | 0.914 | 54 (19.0%) | 0 | 0 |
| PsychChip | 263 | 0 | 244 | 19 | 0 | 0.928 | 43 (16.3%) | 0 | 0 |
| Affymetrix | 137 | 0 | - | - | - | - | 0 | 1 | - |
| ATAC-Seq | 295 | 0 | 260 | 28 | 7 | 0.903 | 55 (18.6%) | 3 | 15 |
| RNA-Seq | 426 | 0 | 414 | 3 | 9 | 0.993 | 3 (0.7%) | 0 | 3 |
| Ribo-Seq | 197 | 0 | - | - | - | - | 50 (25.4%) | 4 | - |
| Total | 1603 | 1 | 1174 | 74 | 21 | - | 201 (12.5%) | 8 | 18 |

*Note:* “Number of sex-matched samples” indicates the number of samples with the same reported sex and SNP-inferred sex.

#### Detect and correct data IDs

We calculated genetic relatedness scores among all the data in the six omics types using GCTA[3]. Based on the distribution of genetic relatedness scores, we extracted the highly related data pairs using a threshold of 0.65 (S2 Fig). We identified a total of 1971 matched pairs and 518 mismatched pairs (Fig 3). We also found eight data that were not related to any other data, including three ATAC-Seq data, four Ribo-Seq data, and one Affymetrix data (S4 Table).

**Fig 3.**
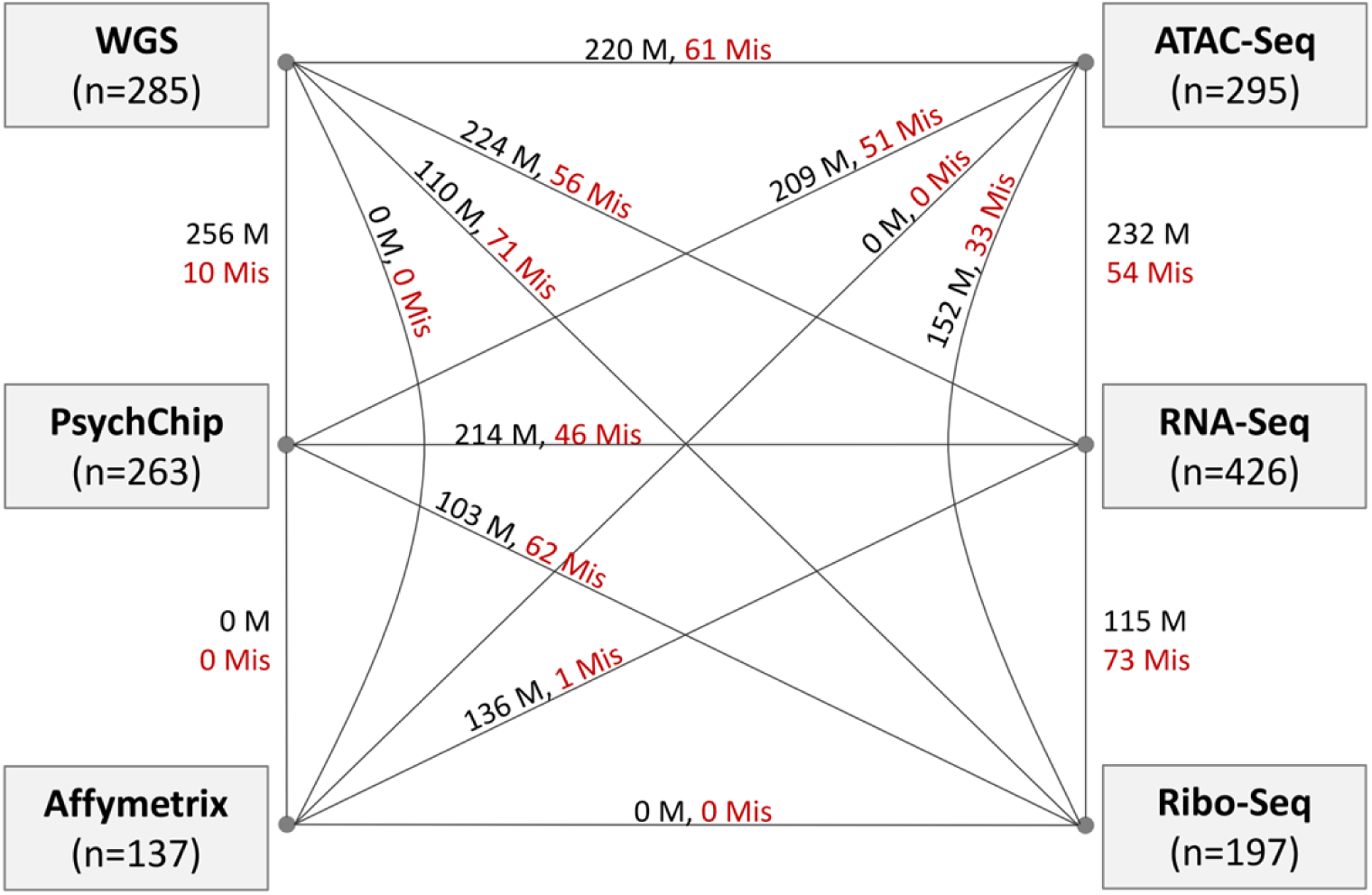
Summary of highly related data pairs. Summary of highly related data pairs among the six omics types. M: Matched pairs, Mis: Mismatched pairs.

Of the 518 mismatched data pairs, 44 pairs have certain switch directions. We used these to train the logistic regression model **(Method)**. For the remaining 404 mismatched data pairs, we used the logistic regression model to predict switch directions and probabilities. Because WGS and PsychChip data were processed with the same sources of DNA, we considered them as one omics type in the regression step. Similarly, we considered ATAC-Seq and Ribo-Seq as one omics type in the regression, as they were processed from the same original tissues. Based on the proportion of samples with concordant SNP-inferred sex and reported sex (Table 1), as well as on our prior knowledge about the data processing of each omics type, we assigned the omics priority as RNA-Seq > Affymetrix > WGS & PsychChip > ATAC-Seq & Ribo-Seq. After using the logistic regression to predict the switch directions and probabilities for the 404 mismatched data pairs, we connected all highly related data pairs to create clusters of paired data. Then, we used the modified topological sorting algorithm to sort the nodes in each cluster and picked the node with the highest score as the final ID for all data in each cluster. In the end, we corrected 201 (12.5%) IDs for data of the six omics types (Table 1, S5 Table).

After correcting data IDs, eighteen data still have mismatched SNP-inferred sexes and reported sexes, including 15 ATAC-Seq data and three RNA-Seq data (S3 Fig). For ATAC-Seq, mismatches are not unexpected since accurately inferring genetics-based sex for some samples is difficult due to their low sequencing coverage in X and Y chromosomes. For RNA-Seq, we found that the XIST-inferred sexes were all consistent with reported sexes for all the samples (S4 Fig), indicating that all the samples in RNA-Seq might have been assigned with their true IDs. As we found three samples with mismatched SNP-inferred sexes and reported sexes, we inferred that it may be inaccurate to estimate genetics-based sexes based on the genotypes called from RNA-Seq data. For Ribo-Seq, both XIST-inferred sexes and F-values reported sexes were more inconsistent than RNA-Seq or DNA-based data. This may suggest that neither sex chromosome genotypes nor XIST gene expression works well to infer genetics-based sexes for Ribo-Seq data.

#### Validate data ID corrections by race group assignment

To confirm that the 201 data IDs were correctly assigned, we used race as an independent validation. We performed Principal Component Analysis (PCA) on samples of the four major racial groups, European, Asian, African, and African American from the 1000 Genomes Project (1000G)[8] (S2 Table) and our BrainGVEX samples. PCA plotted the 1000G and BrainGVEX samples into four racial clusters. Before correcting data IDs, twenty-one samples clustered into wrong racial groups. After correcting data IDs, all samples have concordant races with 1000G samples (S5 Table). For WGS, PsychChip, ATAC-Seq, and Ribo-Seq data that have race-switched data IDs, all were switched back into the correct PCA clusters, indicating that those samples were likely to have been mislabeled and successfully corrected by DRAMS (Fig 4).

**Fig 4.**
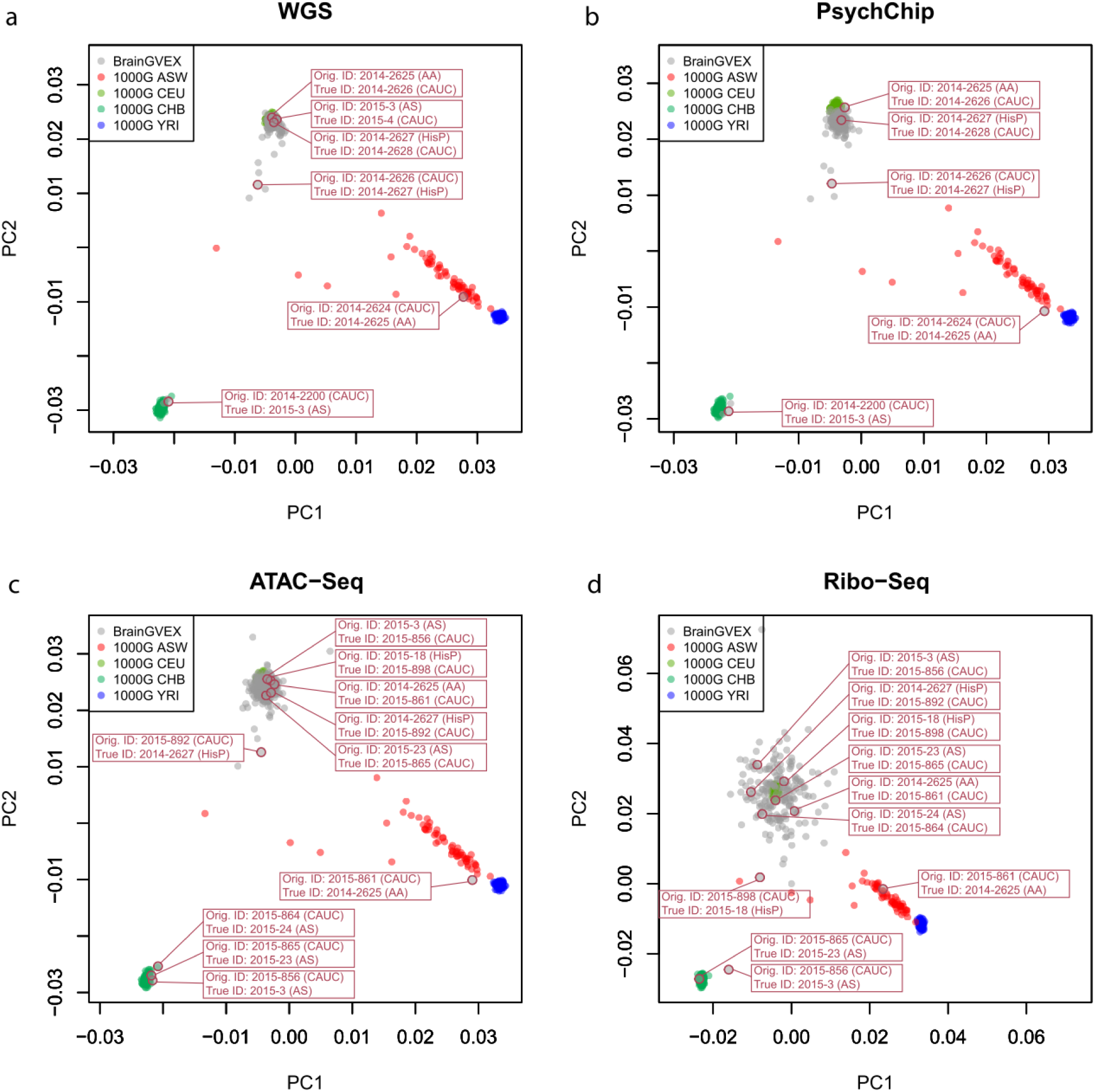
Validation of cross-races switched data. PCA results for BrainGVEX samples (grey dots) and 1000G samples (colored dots) are shown. All data with switched races are marked with their original IDs and new IDs, as well as the corresponding race. The correspondence between BrainGVEX and 1000G races are shown in S2 Table. RNA-Seq and Affymetrix data are not shown as there is no data with switched races. Proportions of switched races data per dataset: WGS: 0.12; PsychChip: 0.12; ATAC-Seq: 0.19; Ribo-Seq: 0.21.

#### Increased number of cis-QTLs after correcting data IDs

We mapped four sets of cis-QTLs based on different data combinations (WGS with RNA-Seq, WGS with Ribo-Seq, PsychChip with RNA-Seq, and PsychChip with Ribo-Seq) for BrainGVEX data before and after correcting data IDs using DRAMS. After correcting data IDs, although the sample sizes were reduced slightly due to the removal of a few unresolved samples, the numbers of cis-QTLs increased by an average of 1.62-fold for the FDR<0.01 cutoff and average 1.54-fold for the FDR<0.05 (Table 2). This relative stability clearly demonstrated the power and importance of correcting data IDs in QTL mapping.

**Table 2.** Increased number of cis-QTLs after correcting data IDs

| QTL type |  | Before<br>correcting IDs | After<br>correcting IDs | Fold |
| --- | --- | --- | --- | --- |
| WGS vs. RNA-Seq | Sample size | 278 | 273 |  |
|  | #QTLs (FDR<0.01) | 57,209 | 96,242 | 1.68 |
|  | #QTLs (FDR<0.05) | 90,231 | 147,942 | 1.64 |
| WGS vs. Ribo-Seq | Sample size | 191 | 187 |  |
|  | #QTLs (FDR<0.01) | 18,178 | 31,345 | 1.72 |
|  | #QTLs (FDR<0.05) | 30,641 | 48,306 | 1.58 |
| PsychChip vs. RNA-Seq | Sample size | 259 | 253 |  |
|  | #QTLs (FDR<0.01) | 48,742 | 76,995 | 1.58 |
|  | #QTLs (FDR<0.05) | 77,925 | 117,711 | 1.51 |
| PsychChip vs. Ribo-Seq | Sample size | 177 | 172 |  |
|  | #QTLs (FDR<0.01) | 15,028 | 20,447 | 1.36 |
|  | #QTLs (FDR<0.05) | 26,350 | 32,209 | 1.22 |
| Total | #QTLs (FDR<0.01) | 139,157 | 225,029 | 1.62 |
|  | #QTLs (FDR<0.05) | 225,147 | 346,168 | 1.54 |

### Sensitivity and specificity in extracting highly related data pairs

To determine the minimum number of SNPs needed to accurately identify highly related data pairs from all random pairs, we randomly selected eight subsets of SNPs numbered from 200 to 10,000 from the BrainGVEX data and re-calculated pair-wise genetic relatedness scores. Five types of omics data were used for this estimation, including WGS, PsychChip, ATAC-Seq, RNA-Seq, and Ribo-Seq. Only the samples with fully matched data IDs among all omics types were used.

Since both the WGS and the PsychChip platforms cover most of the common SNPs in the whole genome, we were able to successfully identify all the highly related data pairs without any false positives, using as few as 200 common SNPs (MAF > 0.1) (S6 Table, S5 Fig). When comparing DNA-based genotypes with data from other platforms, such as RNA-Seq, Ribo-Seq, or ATAC-Seq, we could capture only a small proportion of genotyped loci ranging from 74% to 85%, due to the platforms’ limited coverage of genomic regions. To ensure that enough genotypes are compared, at least 1000 SNP loci should be used to estimate genetic relatedness.

The comparison of data from RNA-Seq, Ribo-Seq, or ATAC-Seq can be problematic since each platform has somewhat different priorities for capturing various genomic regions. Because of this, we found that fewer genotyped loci could be compared, undoubtedly reducing sensitivity and specificity. For example, for the comparison between RNA-Seq and ATAC-Seq data, the proportion of shared SNPs range from 51% to 55%. Even when 1000 common SNPs were used, nine false positive pairs and three false negative pairs were still found (sensitivity: 0.9796; specificity: 0.9996). We recommend using 2000 or more common SNPs to estimate genetic relatedness.

We also did the same analysis using rare SNVs (MAF < 0.1). Rare SNVs are less powerful to distinguish highly related data pairs from random pairs than common SNPs (S7 Table, S6 Fig). It is difficult to identify all highly related data pairs even when 10,000 rare SNVs were used. However, rare SNVs have good specificity. When 1000 rare SNVs were used, only two false positives for WGS versus ATAC-Seq (specificity: 0.9999), and five false positives for RNA-Seq versus ATAC-Seq (specificity: 0.9998) were found. No false positives were found for all the other comparisons.

## Discussion

To meet the demand for reducing the influence of sample mix-ups on multi-omics integrative studies, we developed a tool we refer to as DRAMS to detect and correct mixed-up data IDs. The principle of DRAMS is that genotypes of all omics data assayed on the same individuals should be identical. We directly call genotypes and estimate pair-wise genetic relatedness by calculating the genotype concordance rates among all data to be checked. Therefore, any omics type, as long as it contains genotype information, could be used for DRAMS correction.

Having more omics types bolstered the information available to unlock the potential IDs; however, the increase also results in greater complexity. DRAMS used a logistic regression model followed by a modified topological network sorting algorithm to systematically integrate the genetic relationships, sex concordance, and omics priority to determine the potential ID for each data. The tool performs well in both simulation data and BrainGVEX data. With this design, DRAMS can be applied to an unlimited number of omics data. According to our simulation data, DRAMS performs better as more omics types are included. This is a major advancement of our framework that outperformed existing tools.

Since sex plays such an essential role in verifying data ID, we naturally chose to employ sex information in the design of DRAMS to increase reliability. When applying DRAMS to BrainGVEX data, we used two strategies; we inferred genetics-based sexes from sex chromosome genotypes (SNP-inferred sex) and alternatively, from XIST gene expression (XIST-inferred sex). When applying DRAMS to RNA-Seq data, all data had consistent XIST-inferred sexes and reported sexes after correcting the data IDs. However, three out of 426 data had inconsistent SNP-inferred sexes and reported sexes. For Ribo-Seq data, as XIST is a non-coding RNA, it’s not accurate to estimate genetics-based sex according to XIST expression. Also, due to the low coverage for Ribo-Seq data, estimating genetics-based sex according to sex chromosome genotypes is not confident. Therefore, it is reasonable that we did not see an obvious consistency among XIST-inferred sexes, SNP-inferred sexes, and reported sexes.

We used ethnic information as a validation step for BrainGVEX data ID correction. After correcting data IDs, all of the samples clustered into the correct race with other 1000G reference samples. Although matched race does not necessarily mean correct data ID, it is strong evidence to prove that the data IDs were switched to the correct directions for the mismatched data pairs.

We identified more QTLs after data ID correction. Although this is not direct proof for correct data ID assignment either, it is likely the results of better alignment of different omics types, and consequently largely increased statistical power for QTL analyses.

We used the BrainGVEX data to assess the minimum number of SNPs needed to identify highly related data pairs. We found that common SNPs have greater power in distinguishing highly related data pairs from random pairs. Nonetheless, rare SNVs are also useful as they are unlikely to produce false positive findings. In addition, we also found that when comparing data from different platforms (which cover different genomic regions), a smaller proportion of genotyped loci could be compared between different platforms, indicating that additional SNPs should be used. Based on the BrainGVEX data, we recommend using 2000 or more common SNPs to extract highly related data pairs for most platforms. Nonetheless, since the genotype qualities of different platforms may differ substantially, we recommend including as many variants as possible, even rare variants, to fortify the sensitivity and specificity of the DRAMS tool.

Matching omics data is only the first step in the process. Assigning the correct data ID, typically associated with sample demographic information and phenotypic data, is another important step. Conceivably, some analyses are more sensitive to sample information, covariates (sex, diagnosis, etc.) than others. Mis-assigned sample information severely affects some analyses, such as differential gene expression and case-control comparison. DRAMS can correct mix-ups and identify correct labels and associated sample information, before conducting the integrative analyses.

Currently, only a few projects have produced multi-omics data, like the data generated in Drs. Gilad and Pritchard’s lab[9], data from the Genotype-Tissue Expression (GTEx) project [10], and ROSMAP[11, 12]. As more large-scale multi-omics data will be generated in the near future to address problems of regulatory networks and causal relationships, we expect that DRAMS will be helpful for those studies.

## Materials and Methods

### Sample resources

A total of 440 individuals from the PsychENCODE BrainGVEX study[6] with six types of omics data were used to validate data IDs and assign the potential IDs. BrainGVEX was part of the PsychENCODE project focusing on gene expression regulation in human brain FC region (Frontal Cortex). The samples include 420 Caucasians, 2 Hispanic, 1 African American, 3 Asian American, and 14 unknown (S1 Table). The six omics types included 1) 285 samples (176 males, 106 females, and 3 unknown-sex samples) of low-depth Whole Genome Sequencing (WGS, average depth: 5×) data; 2) 426 samples (274 males, 152 females) of RNA-Seq data; 3) 295 samples (180 males, 112 females, and 3 unknown-sex samples) of Assay for Transposase-Accessible Chromatin using Sequencing (ATAC-Seq) data,; 4) 197 samples (122 males, 70 females, and 5 unknown-sex samples) of Ribosome Sequencing (Ribo-Seq) data; and, SNP array data from two platforms, including 5) 137 samples (92 males, 45 females) of Affymetrix 5.0 450K (Affymetrix) data; and, 6) 263 samples (163 males, 100 females) of Psych v1.1 beadchips (PsychChip) data.

### Genotype calling from data of each omics type

We used the same pipeline to call genotypes for all sequencing data. For each dataset in FASTQ format, all reads were mapped to the human reference genome (hg19) using BWA [13] after sequencing adapters and low-quality bases were removed. PCR duplications were removed using the MarkDuplicates package in Picard tools (http://broadinstitute.github.io/picard/). Then, GATK IndelRealigner and BaseRecalibrator were used to recalibrate the mapping quality of the reads [14]. For each omics type, genotypes were called using GATK HaplotypeCaller for all samples jointly. Each set of omics data were processed separately.

### Estimation of sample contamination

Two methods were used to check sample contamination. One is VerifyBamID[7], which because it requires both BAM files and VCF files as input, can only be applied to sequencing data. The results include a parameter “FREEMIX” (0-1 scale), which indicates the proportion of non-reference bases observed in reference sites. This parameter can be used as an indicator of sample contamination. As an alternate method, we wrote a Linux script to directly calculate the heterozygous rate based on genotypes. This ran faster than the VerifyBamID approach. We defined the samples with FREEMIX >0.3 in VerifyBamID and defined heterozygous rates >0.3 as contaminated samples, which were removed from subsequent analyses. Only heterozygous rates were calculated for PsychChip and Affymetrix samples, as they are not supported by VerifyBamID.

### Infer genetics-based sexes

We used the Plink software (“--check-sex” module in “ycount” mode) to calculate F-values for data of WGS, PsychChip, ATAC-Seq, RNA-Seq, and Ribo-Seq [15]. This method is mainly based on the X chromosome heterozygosity. It also uses the Y chromosome call rate to improve the accuracy of sex estimates. Basically, the F-values were in bi-modal distribution. Based on the distribution, we were able to select a threshold for each omics type and infer sexes (SNP-inferred sexes) for each data. The data with F-value larger than the threshold were classified as males, while the others were classified as females. We did not infer SNP-inferred sexes for Ribo-Seq data since an obvious bi-modal distribution of F-values was not apparent.

For RNA-Seq and Ribo-Seq data, we also inferred sexes based on XIST (X-inactive specific transcript) gene expression levels (XIST-inferred sex). XIST is a noncoding RNA that is only expressed in cells containing at least two X chromosomes [16]. Normally, the XIST gene is only expressed in female samples. In this study, we considered samples with XIST expression larger than 2 (TPM, Transcripts Per Kilobase Million) as females.

We compared the reported sex and genetics-based sex for each data in each omics type and calculated the sex concordance rate for each omics type. Samples with unknown SNP-inferred sexes were not included when calculating the sex concordance rate. The sex concordance rate can represent a parameter indicating omics priority in the logistic regression model (See “Estimate switch directions and probabilities for mismatched data pairs”).

### Estimate genetic relatedness and extract highly-related data pairs

Two tools were used to estimate genetic relatedness among data of multiple omics types and extract highly-related data pairs: GCTA[3] and NGSCheckMate[4]. For GCTA, GRM module was used. Basically, the genetic relatedness scores are distributed bimodally, so that one peak with higher scores indicates highly-related data pairs while the other peak indicates random (unrelated) data pairs. In this way, one can “eyeball” the distribution to determine the thresholds of genetic relatedness scores between every two omics datasets (Fig 1). Using that method, we applied a threshold of 0.65 for our BrainGVEX data and extracted highly related data pairs in different omics types. For NGSCheckMate, we ran the software in VCF mode with an “-f” parameter to enact a strict VAF correlation filter. We calculated the concordance rates of the two tools based on BrainGVEX data.

### Sensitivity and specificity in extracting highly related data pairs

To determine the minimum number of variants required to extract highly related data pairs from all combination of data pairs, we re-calculated genetic relatedness scores using the BrainGVEX data based on subsampled SNPs. Data from five omics types were used, including WGS, PsychChip, ATAC-Seq, RNA-Seq, and Ribo-Seq. We used only the samples that fully matched in all the five omics types. For comparisons between each two omics types, we randomly selected 200, 400, 600, 800, 1000, 2000, 5000, and 10,000 SNPs that were called by both omics types and calculated pair-wise genetic relatedness scores using GCTA. The data pairs with genetic relatedness scores larger than 0.65 were classified as *highly related data pairs* and those pair with smaller scores were classified as *unrelated data pairs*. For each comparison, we calculated the true positive rate and false negative rate. We did the same analysis for common (MAF>0.1) and for rare (MAF<0.1) variants.

### Estimate switch directions and probabilities for mismatched data pairs

To estimate the possible switch directions and the probabilities for a mismatched data pair, the key point must be focusing on determining which data are more likely to bear the true ID. We used a logistic regression model (Formula 1) to compare the two data presented in each mismatched data pair and to estimate the possible switch direction and its probability (Fig 1). Assuming the mismatched data pair “A” and “B”, if “B” is more likely to have the true ID, then the appropriate switch direction is from “A” to “B”. Three parameters (*x*_*1*_, *x*_*2*_, and *x*_*3*_, values range from 0 to 1) were used in this model. The first parameter, *x*_*1*_, indicates which data matched with more data in other omics types (Formula 2). The second parameter, *x*_*2*_, indicates which data are more likely to have correct reported sex (Formula 3); and the third, *x*_*3*_, is a user-defined parameter indicating the rank of omics priority. If two or more omics types were processed under the same condition or process, these omics types may have the same sample mis-labeling, a user could combine these omics types into one type in the regression. If *x*_*3*_ is not specified, the rank of omics priority will be defined based on the proportion of data that have reported sex and genetics-based sex-matched for each omics type (Formula 4).

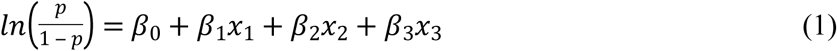

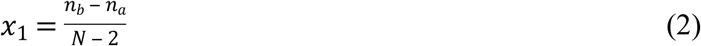

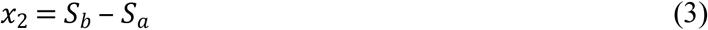

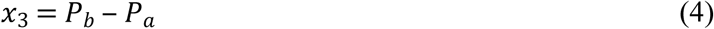

In formula 2, N is the total number of omics types; and correspondingly, n_a_ and n_b_ are the number of matched data from other omics types for data “A” and “B”, respectively. In formula 3, S_a_ and S_b_ indicate the sex matching level for data “A” and “B”, respectively. Taking S_a_ as an example, the original value of S_a_ would be 0. If the reported sex and genetics-based sex in data “A” are matched, S_a_ would be plus 0.5. If the reported sex of data “A” and the genetics-based sex of data “B” are matched, which means that the reported sex and genetics-based sex are matched after switch ID from “B” to “A”, then S_a_ would be plus 0.5. In formula 4, P_a_ and P_b_ represent the proportion of data with matched reported sex and genetics-based sex for data “A” and “B”, respectively.

A set of hand-picked high-confidence mismatched data pairs with well-defined switch directions were used as a training set for the logistic regression model. The directions of the high-confidence mismatched data pairs were determined based on sample relationships and sex matching. We connected all the highly related data pairs among multiple omics types and created multiple clusters. We extracted the high-confidence mismatched data pairs based on the following conditions: 1) If only one data had a different ID, which needs to be corrected, from the others in the cluster; 2) If the reported sex and genetics-based sex were matched after correcting that ID based on other data in the cluster.

After training, the values for β_0_, β_1_, β_2_, and β_3_ were defined. Then, the model was used to predict switch directions and probabilities for the rest of the data. For the results, if p > 0.5, the switch direction will be from “A” to “B”; conversely, if p < 0.5, the switch direction will be from “B” to “A”; and, if p = 0.5, the switch direction will be uncertain. The values of p for both directions (p > 0.5, p < 0.5) were normalized (with a range from 0 to 1 respectively) to indicate the probabilities of switch directions.

### A modified topological sorting algorithm to determine the potential IDs

We connected all highly related data pairs and generated multiple clusters. In each cluster, each data was represented by a node, and each data pair was represented by an edge. Based on the logistic regression model results, the switch direction and probability for each mismatched data pair corresponded to the direction and weight of each edge. For each matched data pair, we did not assign a direction or weight for the edge. Since all the data presented in a cluster were supposed to have one unique ID, we used a modified topological sorting algorithm to sort all nodes in each cluster to determine the potential ID for each data.

The modified topological sorting algorithm was based on the indegrees and outdegrees weighted by the switch probabilities which had been calculated in the logistic regression model. For each node in each cluster, we calculated the difference between weighted indegree and weighted outdegree. Afterward, we sorted the nodes in each cluster based on the difference between weighted indegrees and outdegrees. For each cluster, we used the data ID with the highest priority as the final ID for all data presented in the cluster.

### Simulation data

We generated simulation data to test the performance of DRAMS on correcting data IDs. A range of sample sizes (50, 100, 150, 200, 250, and 300) was used, with each dataset having 50% females and 50% males. For each individual, we generated data of multiple omics types ranging from three to six and assigned IDs for each data. The IDs of different omics types for the same individual were pair-wise connected as highly related data pairs. To simulate sample mix-ups, we randomly shuffled the following gradient proportions of data IDs for each omics type: 5%, 10%, 15%, 20%, 25%, and 30%. In total, we generated 144 simulated datasets (six sets of sample size × four sets of omics type × six proportions of mixed-up sets). In an attempt to mimic reality, we randomly introduced mislabeled sexes for 2% of samples in our simulation data. In addition, since samples are often swapped within the same batch in reality, we divided the samples into several batches, with each batch containing 25 samples. The data ID shuffling all occurred within batches. For each set of simulation data, we corrected data IDs using DRAMS and calculated the proportion of mix-ups being successfully corrected. We simulated the process 100 times for each dataset.

### Validate data ID corrections by racial assignment

To validate the tool, we started by using genotypes that shared loci in BrainGVEX and the 1000 Genomes project (1000G)[8]. For 1000G, we used only the following samples: ASW (African Ancestry in Southwest US), CEU (Utah Residents with Northern and Western European Ancestry), CHB (Han Chinese in Beijing, China), and YRI (Yoruba in Ibadan, Nigeria). We first removed genotypes with MAF < 0.01. Then, we used GCTA[3] to estimate genetic relationships among individuals and performed PCA analysis for both BrainGVEX and 1000G samples jointly. We used samples from 1000G that were located in the same cluster as the reference, after which we compared the races of samples from BrainGVEX with the reference before and after correcting data IDs. Since BrainGVEX and 1000G used different nomenclatures for races, we used analogical names or close races between BrainGVEX and 1000G to align the data (S2 Table).

### Detect QTLs before and after correcting mix-ups

We used FastQTL[17] to map QTL within the BrainGVEX samples both before and after correcting data IDs. We defined the cis-QTL region as 1 million base pairs between the SNP marker and the gene body. Since we intend to test whether the number of cis-QTLs increased, only chromosome 1 was used to save computing time. R package “qvalue”[18] was used for multiple tests. We tested four types of cis-QTLs calculations: WGS with RNA-Seq, WGS with Ribo-Seq, PsychChip with RNA-Seq, and PsychChip with Ribo-Seq. For RNA-Seq and Ribo-Seq samples, we used log2 transformed CPM quantification data calculated by VOOM[19]. We selected 30 hidden factors as covariates using the PEER software[20] for RNA-Seq and Ribo-Seq samples. We used two cutoffs (FDR < 0.05, FDR < 0.01) for the cis-QTL results.

### Code availability

The code of DRAMS was implemented in Python3 and deposited in GitHub (https://github.com/Yi-Jiang/DRAMS). We released the data preprocessing codes, including genotype calling, sample contamination checking, genetics-based sex inference, genetic relatedness score calculation, and extracting highly related data pairs. We also provided a guideline for using Cytoscape to visualize sample relationships within networks [21].

## Author’s contributions

YJ developed the tool, tested the tool on both simulation data and BrainGVEX data, and led the manuscript writing. GG, KG, and AWS generated the data and helped in dealing with BrainGVEX data. KPW participated in the data production. YX, LH, QWang, QWei, RC, and SL helped in developing the tool. YX revised the manuscript. CL, CC, and BL supervised the overall study design, guided analyses, and revised the manuscript. All authors read and approved the final manuscript.

## Competing interests

The authors have declared no competing interests.

## Acknowledgements

This work was supported by National Natural Science Foundation of China [31571312 and 81401114 to C.C., 31871276 to C.L.], the National Key Plan for Scientific Research and Development of China [2016YFC1306000 to C.C.], and National Institutes of Health [U01 MH103340-01 and 1R01ES024988 to C.L.]. Thanks the China Scholarship Council (CSC) for financial support. Thanks Liz Kuney and Richard Kopp for revising the manuscript. Thanks Hai Yang and Ying Ji for giving advice on presenting the work.

